# Yeast eukaryotic initiation factor 4B remodels the mRNA entry site on the small ribosomal subunit

**DOI:** 10.1101/2023.01.18.524494

**Authors:** Ayushi Datey, Faisal Tarique Khaja, Huma Rahil, Tanweer Hussain

## Abstract

Eukaryotic initiation factor 4B (eIF4B) belongs to the eIF4 group of factors that help in mRNA recruitment to the ribosomal preinitiation complex (PIC) in all eukaryotic organisms. eIF4B stimulates the helicase activity of eIF4A and helps in the formation of the 48S PIC by facilitating mRNA recruitment. However, there is no clear understanding of the location of eIF4B on the 40S and how eIF4B helps in the recruitment of mRNAs. In this work using cryo-electron microscopy, we show that yeast eIF4B binds to the 40S ribosomal subunit at the mRNA entry channel making contacts with ribosomal proteins uS10, uS3, and eS10 and ribosomal rRNA helix h16. The yeast eIF4B position on the 40S overlaps with the RRM domain of eIF3g indicating that the binding of eIF4B may trigger the relocation of the eIF3 b-g-i module to the subunit interface. The 40S head is in partially open conformation that may facilitate the release of eIF3j and hence aid mRNA recruitment and scanning. The structural analysis of yeast eIF4B-bound ribosomal complex provides insight into possible events during mRNA recruitment.

## Introduction

Eukaryotic translation initiation is a highly intricate process requiring a myriad of initiation factors (Aitken and Lorsch, 2012). Assembly of the initiation factors eIF1, eIF1A, eIF3, eIF5 and the eIF2-GTP-initiator tRNA ternary complex (TC) on the 40S ribosomal subunit to form the 43S pre-initiation complex (PIC) marks the commencement of initiation (Hinnebusch, 2017). The eIF4F complex binds and activates the mRNA for initiation and recruits it to the 43S PIC, resulting in the formation of the 48S PIC. This eIF4F complex is composed of the mRNA 5ʹ cap-interacting protein eIF4E, the ATP-dependent RNA helicase eIF4A and the scaffold-providing eIF4G protein (Merrick, 2015; Mishra *et al*., 2020). The ATPase and helicase activity of eIF4A is enhanced by eIF4G together with the initiation factor eIF4B (Andreou and Klostermeier, 2014).

Among all the stages of translation initiation, recruitment of the mRNA on the 43S PIC forms the rate-limiting step (Fraser, 2015; Aylett and Ban 2017). As mentioned above, preparation or activation of the mRNA for initiation requires the assembly of the eIF4F complex on its 5ʹ cap. The presence of secondary structural elements in the 5ʹ UTR hinders initiation (Jackson *et al*., 2010; Kozak, 1986; Uppala *et al*., 2022). For this, the factor eIF4A, an ATP-dependent RNA helicase, is essential as it melts the secondary structures in the mRNA 5ʹ UTR (Rogers *et al*., 1999), thus allowing the efficient formation of the 48S PIC. The helicase activity of eIF4A is enhanced in the presence of factors eIF4G and eIF4B (Andreou and Klostermeier, 2014; Rogers *et al*., 1999). In yeast, eIF4B interacts directly with the 40S ribosomal protein rps20 (uS10) located in the head region of the ribosome, thus helping in the recruitment process (Walker *et al*., 2013). The mRNAs harbouring long and extensively structured 5ʹ UTR also require eIF4B for their efficient translation initiation to occur. However, this requirement is not based on the eIF4A-helicase activity-enhancing role of eIF4B. Recent reports have also indicated the presence of a class of mRNAs that are hyper-dependent on eIF4B while exhibiting less dependence on eIF4G and/or eIF4A (Liu *et al*., 2021; Sen *et al*., 2016). The ability of eIF4B to directly interact and bind the 40S ribosomal subunit seems to be crucial for the translation of these mRNAs. Therefore, it appears that eIF4B also performs a function independent of eIF4A/4F.

Polysome profiling experiments have also reported the importance of eIF4B in translation initiation as there is a significant decrease in polysome to monosome ratio in the absence of eIF4B (Walker *et al*., 2013). Among all the eIF4 groups of factors from different eukaryotes, the characterization of eIF4B has been most difficult owing to the lack of sequence similarity. However, eIF4B has been characterized from organisms like *Saccharomyces cerevisiae, Homo sapiens, Drosophila melanogaster*, and *Arabidopsis thaliana* and this has helped identify putative eIF4B encoding genes in other organisms. eIF4B from the fungi, plants, and mammals form three separate groups in phylogenetic tree analysis (Hernandez and Vazquez-Pianzola, 2005). Even though eIF4B is the least conserved eIF4 factor, the function of eIF4B in enhancing the recruitment of mRNAs seems to be conserved from yeast to plants to humans (Mayberry *et al*., 2009; Sen *et al*., 2016; Shahbazian *et al*., 2010).

Yeast eIF4B has four putative functional domains: an N-terminal domain (NTD), an RNA Recognition Motif (RRM), a 7-repeats region (7-RR) made of 7 imperfect repeats of 26 amino acid residues, and a C-terminal domain (CTD) (Zhou *et al*., 2014). Among all of them, RRM is the only structured domain, while the rest are predicted to be disordered (Mishra *et al*., 2020). Also, RRM is the only conserved domain among yeast and human eIF4B. However, certain conserved motifs are present in the NTD, and a region homologous to the core of the yeast 7-RR is present in human eIF4B after the RRM domain and just before the DRYG repeat domain (Zhou *et al*., 2014). In yeast, the NTD and 7-RR have been reported to be responsible for the interaction of eIF4B with 40S, independent of the RRM domain (Walker *et al*., 2013). It has been suggested that the binding of eIF4B to the 40S ribosomal subunit may promote conformational changes in the mRNA binding channel of the ribosome, making it more accessible (Liu *et al*., 2021). However, there is no structural understanding of the conformational changes in the 40S upon eIF4B binding or how eIF4B helps in the recruitment of mRNAs. Along with this, there is no clear understanding of the location of eIF4B on the 40S as well as the points of contact eIF4B makes with the ribosome. The knockout of eIF4B in yeast leads to growth defects whereas its knockdown in human cells leads to decreased protein synthesis (Altmann *et al*., 1993; Shahbazian *et al*., 2010). These studies suggest that the role played by eIF4B in translation initiation is crucial for protein synthesis and more importantly the interaction of eIF4B with the 40S is critical. In this study, we aim to understand the association of 40S with eIF4B and how this interaction helps in translation initiation.

In this work, we reconstructed the cryo-electron microscopy (cryo-EM) map of the 40S bound with eIF4B and show that eIF4B binds to the yeast 40S ribosomal subunit at the mRNA entry channel on its solvent-exposed side (Figure 1A and 1B). The 40S-eIF4B map provides information about the interacting partners of eIF4B along with suggesting the possible ways in which eIF4B helps in mRNA recruitment.

**Figure 1:**
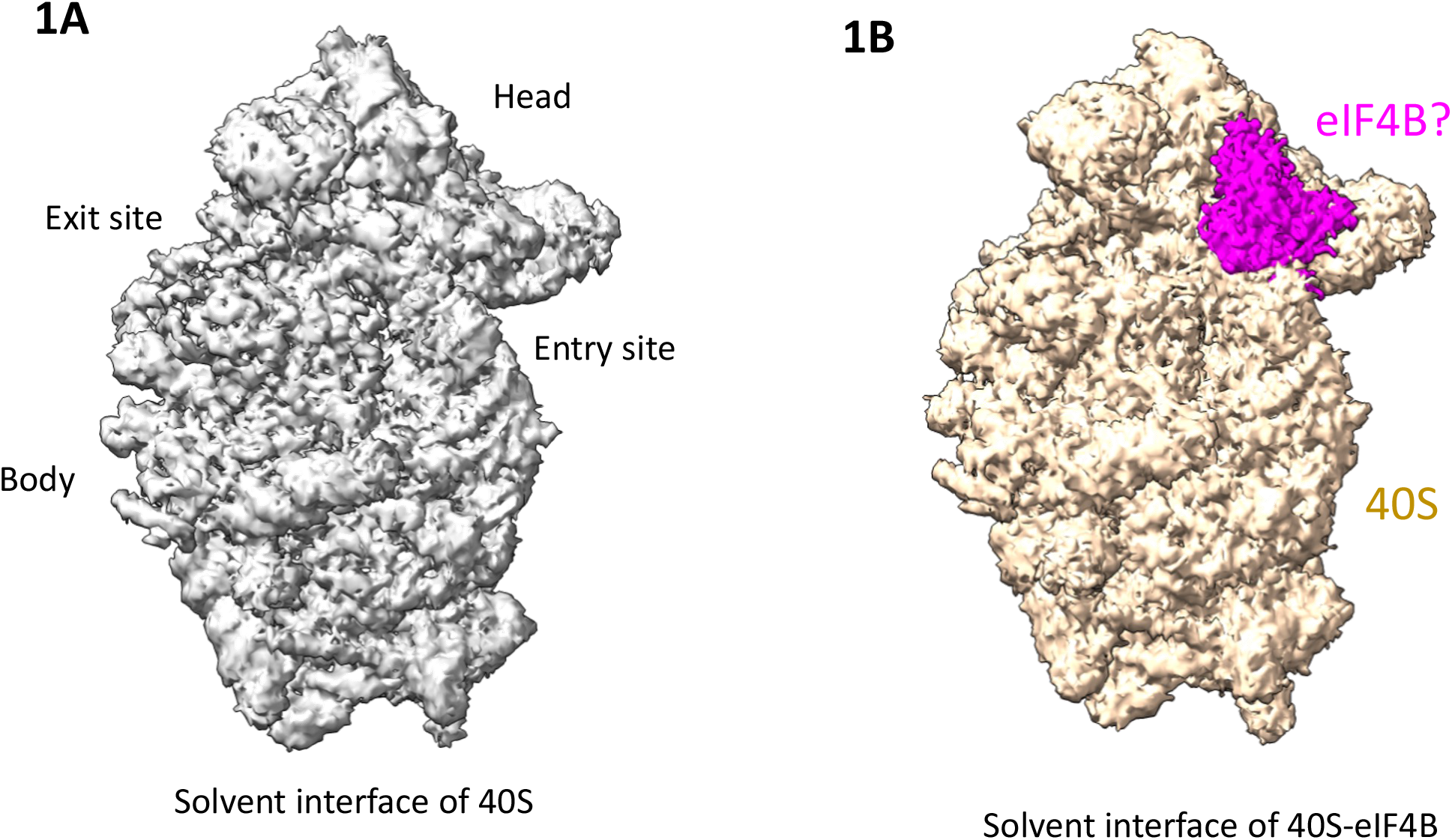
Cryo-EM maps of empty and eIF4B-bound small ribosomal subunits. (A) Cryo-EM map of empty 40S or apo-40S. (B) Cryo-EM map of 40S-eIF4B with the extra density tentatively assigned to eIF4B highlighted in magenta.

## Results and Discussion

### Position of yeast eIF4B on 40S ribosomal subunit

To elucidate the interaction of the 40S with eIF4B, we reconstituted the complex using purified, recombinant *Saccharomyces cerevisiae* eIF4B and purified *Kluyveromyces lactis* 40S. We obtained two maps (Figures S1 and S2), one of the empty 40S (Figure 1A) and another one of 40S-eIF4B (Figure 1B, Table 1, Movie 1) at moderate resolutions of 4.0 Å and 4.6 Å, respectively during the cryo-EM data processing. The 40S map was an extra density was observed at the mRNA entry channel of the 40S on the solvent-exposed side (Figure 1B, Movie 1) in the map obtained from the remaining 25% of the particles. We have tentatively assigned this density to eIF4B; therefore, this map is referred to as the 40S-eIF4B map.

**Table 1:** Refinement and model statistics.

| **Data Collection** | **40S-eIF4B** |
| --- | --- |
| Particles | 593,832 |
| Pixel Size (Å) | 1.17 |
| Voltage (kV) | 200 |
| Detector | K2 |
| Electron Dose (e-/ Å<sup>2</sup>) | 50 |
| Defocus range | -1.5 to -3 |
| Exposure rate (e-/ Å<sup>2</sup>/frame) | 2.5 |
| Data Collection Software | Latitude-S |
| **Data Processing** |  |
| Software | RELION 3.1.1 |
| Usable Micrographs | 7,975 |
| Particles (40S-eIF4B/40S) | 108,616/465,216 |
| Resolution (Masked 0.143 FSC) 40S-eIF4B/40S (Å) | 4.6/4 |
| **Model Composition** |  |
| Chains | 39 |
| Non-hydrogen atoms | 76,299 |
| Protein residues | 4,874 |
| RNA bases | 1,780 |
| **Refinement (40S-eIF4B)** |  |
| Resolution used for refinement (Å) | 4.6 |
| R.M.S. deviations |  |
| Bonds (Å) | 0.006 |
| Angles (°) | 0.985 |
| d FSC 0.143 (Å) | 4.6 |
| d 99 (Å) | 4.7 |
| d model (Å) | 4.1 |
| d FSC model 0.5 | 4.8 |
| **Validation (proteins)** |  |
| Molprobity score | 2.82 |
| Ramachandran plot (%) |  |
| Favoured | 80.41 |
| Allowed | 19.28 |
| Outliers | 0.31 |
| **Validation (RNA)** |  |
| Correct sugar puckers (%) | 97 |
| Good backbone conformations (%) | 45 |
| **Model vs. Data** |  |
| CC (mask) | 0.72 |
| CC (box) | 0.78 |
| CC (peaks) | 0.66 |
| CC (volume) | 0.71 |

The extra density originates near uS3 and rRNA helix h16 and contacts eS10 and uS10 (Figures 2A, 2B, S3A and S3B). Since the local resolution of the extra density for eIF4B ranges from 8-10 Å (Figure S3C) no model building could not be performed. The volume of this density is big enough to accommodate a protein having a molecular weight of 8-10 kDa. Hence it is likely that either the NTD (10 kDa), a part of the 7-RR (19 kDa) or portions of both NTD and 7-RR could be present here, as these two domains have been shown to interact directly with the 40S in yeast (Walker *et al*., 2013). Only two of these seven internal repeats of the 7-RR are sufficient for the activity of eIF4B with the same expression level as that of the wildtype (Zhou *et al*., 2014).

**Figure 2:**
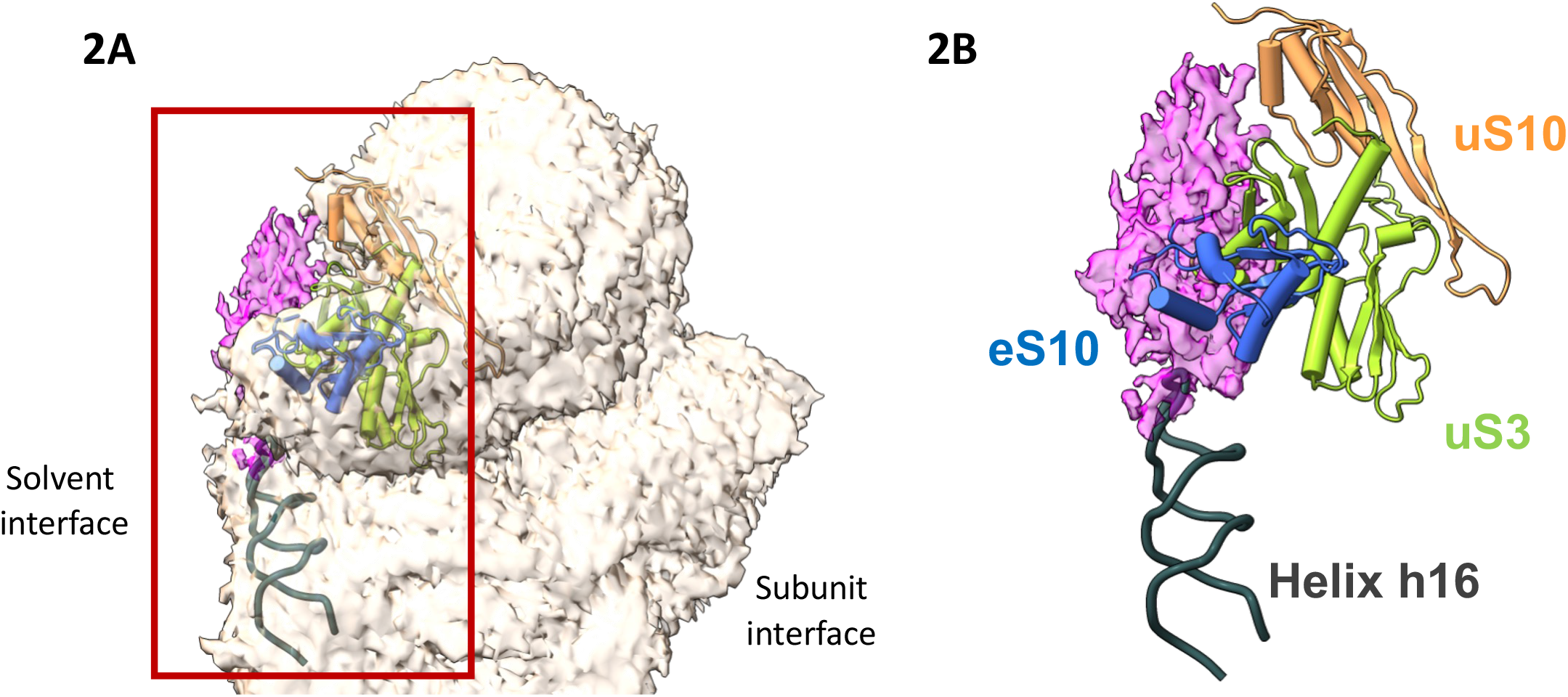
Interacting partners of eIF4B on the 40S. (A) The location of eIF4B on the 40S is on its solvent-exposed side and near the mRNA channel entry. (B) uS3, uS10, eS10 and rRNA helix h16 are the interacting partners of eIF4B on the 40S (uS3 – yellow-green, uS10 – coral, eS10 – blue, h16 - Gray).

eIF4B is reported to interact with uS10 in yeast (Walker *et al*., 2013). Cross-linking and affinity mass spectrometry-based studies have also shown the interaction of yeast eIF4B with both uS3 and eS10 (Rossler *et al*., 2019). A similar study of human eIF4B using affinity capture mass spectrometry showed that it interacts with uS3, uS10, and eS10 (Li *et al*., 2018), suggesting a conserved mode of interaction of eIF4B with the 40S across eukaryotes. A cryo-EM study of partial human 48S PIC had placed the RRM domain of eIF4B near the mRNA entry channel of the ribosome on the inter-subunit interface (Eliseev *et al*., 2018). This placement was supported by the cross-linking mass spectrometry experiment of eIF4B and 40S.

The amino acids present in the interface of human eIF4B-RRM and ribosomal protein uS3 are conserved in the yeast eIF4B-RRM domain and yeast uS3 as well, suggesting that like in humans, yeast eIF4B-RRM domain may occupy this position (Figure 3A and 3B). A comparison of human 48S PIC (Eliseev *et al*., 2018) and 40S-eIF4B map comparison shows that the human eIF4B-RRM occupies the same position as the extra density in the 40S-eIF4B map (Figure 3C and S4A). Although it fits in the density, the RRM domain (9 kDa) was reported to be dispensable for the binding of eIF4B to the 40S in yeast (Walker *et al*., 2013) and the low resolution of the density in 40S-eIF4B map prevents us from assigning any eIF4B domain to this density. It is also interesting to note that the same site is also the binding site for the RRM domain of eIF3g as well in both yeast and humans (Brito Querido *et al*., 2020; Kratzat *et al*., 2021). Although the positions of the RRMs modeled in these PICs (Brito Querido *et al*., 2020; Eliseev *et al*., 2018; Kratzat *et al*., 2021) differ slightly, the main interacting interface is formed by uS3 (Figure 3D). Strikingly, sequence analysis of RRMs of eIF4B and eIF3g from yeast to humans show that the contact points of eIF3g-RRM with uS3 are conserved in eIF3g (Figure 3E) but not in eIF4B (Figure 3F). Thus, further studies are needed to unambiguously model yeast eIF4B-binding region with the 40S. This map shows that yeast eIF4B remodels the mRNA entry site by binding at the pocket formed by uS3, uS10, eS10, and h16. Since eIF4B interacts with eIF4A (Andreou *et al*., 2017), this 40S-eIF4B map indirectly hints at the possible location of eIF4A near the 40S mRNA channel entry to resolve the secondary structure of mRNA before it enters the mRNA channel for translation. The plausible consequences of eIF4B binding to the 40S that aid in mRNA recruitment are discussed below.

**Figure 3:**
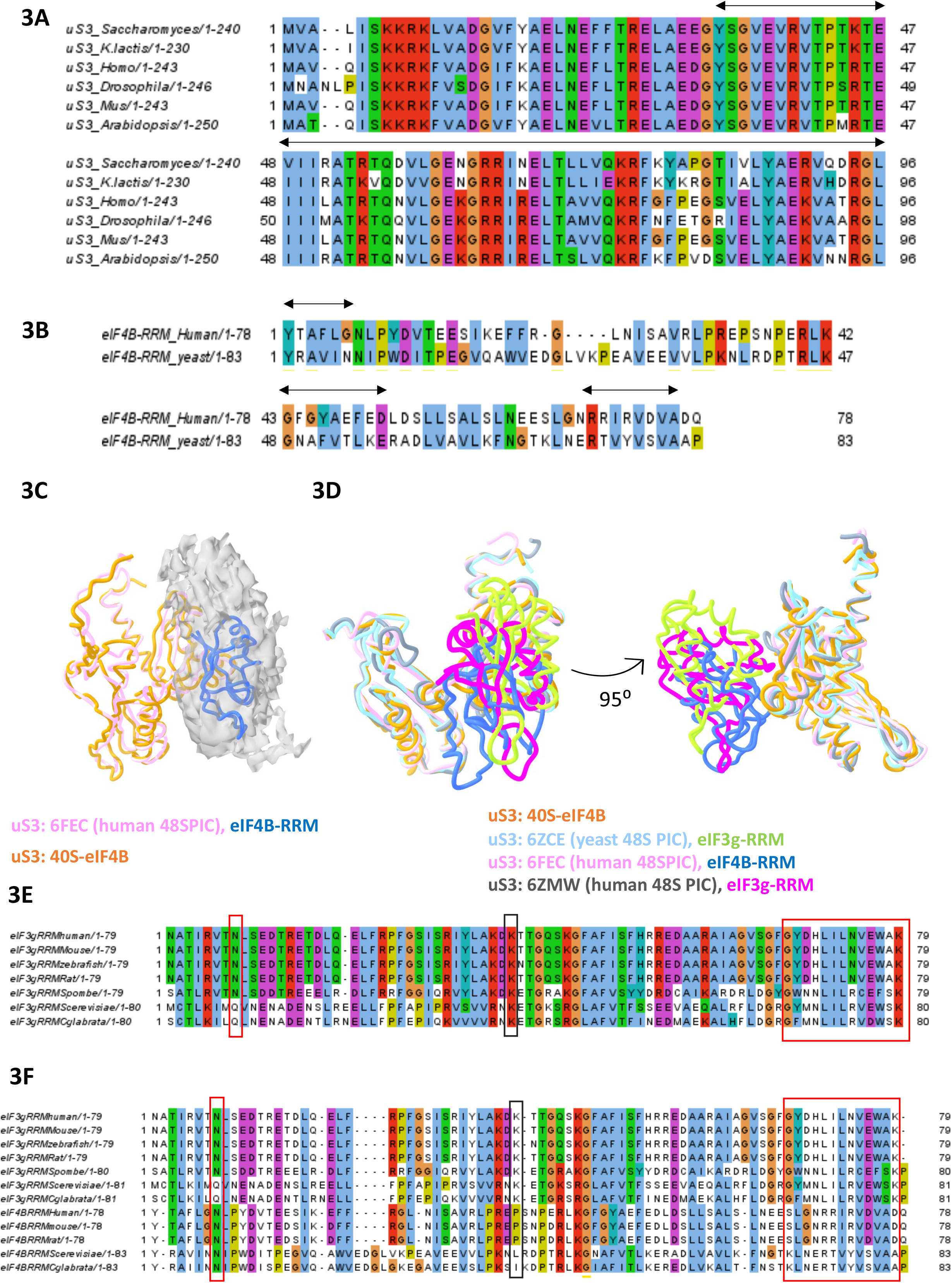
uS3 forms the major interacting interface with the RRM domains of both eIF3g and eIF4B. (A) Multiple sequence alignment of uS3 from eukaryotic model organisms show that the uS3 amino acid residues present at the 40S and eIF4B interacting interface are conserved (marked by black arrows). (B) Sequence alignment of the of human eIF4B-RRM and yeast eIF4B-RRM shows that the eIF4B-RRM residues interacting with the 40S are also conserved in yeast eIF4B (marked by black arrows). (C) eIF4B-RRM from the partial human 48S PIC (PDB: 6FEC) overlaps with the eIF4B density of the yeast 40S-eIF4B map. (D) uS3 forms a common interacting site for the RRM domains of both eIF4B and eIF3g from both humans and yeast. (PDB: 6ZCE-native yeast 43S PIC, 6FEC and 6ZMW-partial human 48S PIC). (E) Multiple sequence alignment of the RRM domain of eIF3g from eukaryotes model organisms show that the amino acid residues interacting with 40S protein uS3 (marked by red boxes) and rRNA helix h16 (dark blue box) are conserved across all organisms. (F) Structure based multiple sequence alignment of RRMs of eIF3g and eIF4B shows that most of residues in RRM of eIF3g that interact with uS3 are not conserved in the RRM domains of eIF4B (red boxes show residues interacting with uS3 and dark blue box shows residue interacting with helix h16).

### Possible consequences of 40S-eIF4B interaction

The mRNA channel of the 40S has a latch formed by helices h18 and h34 present in the body and head of the ribosome, respectively (Figure 4A). This latch needs to be in an open conformation for the mRNA to enter and accommodate in the channel for initiation (Llacer *et* al., 2015). In the 40S-eIF4B map, the latch is partially open suggesting that eIF4B might help stabilize an open state of the small ribosomal subunit (Figure 4). An open state would aid in mRNA recruitment as well as in scanning of 5’UTR as the latch between h18 and h34 is open and the mRNA channel is widened. The binding of eIF1 and eIF1A in the early steps of the canonical initiation pathway opens the mRNA latch (Llacer *et al*., 2015). This study suggests that binding of eIF4B would further stabilize the open form and thus contribute towards mRNA recruitment and scanning of long 5’UTR.

**Figure 4:**
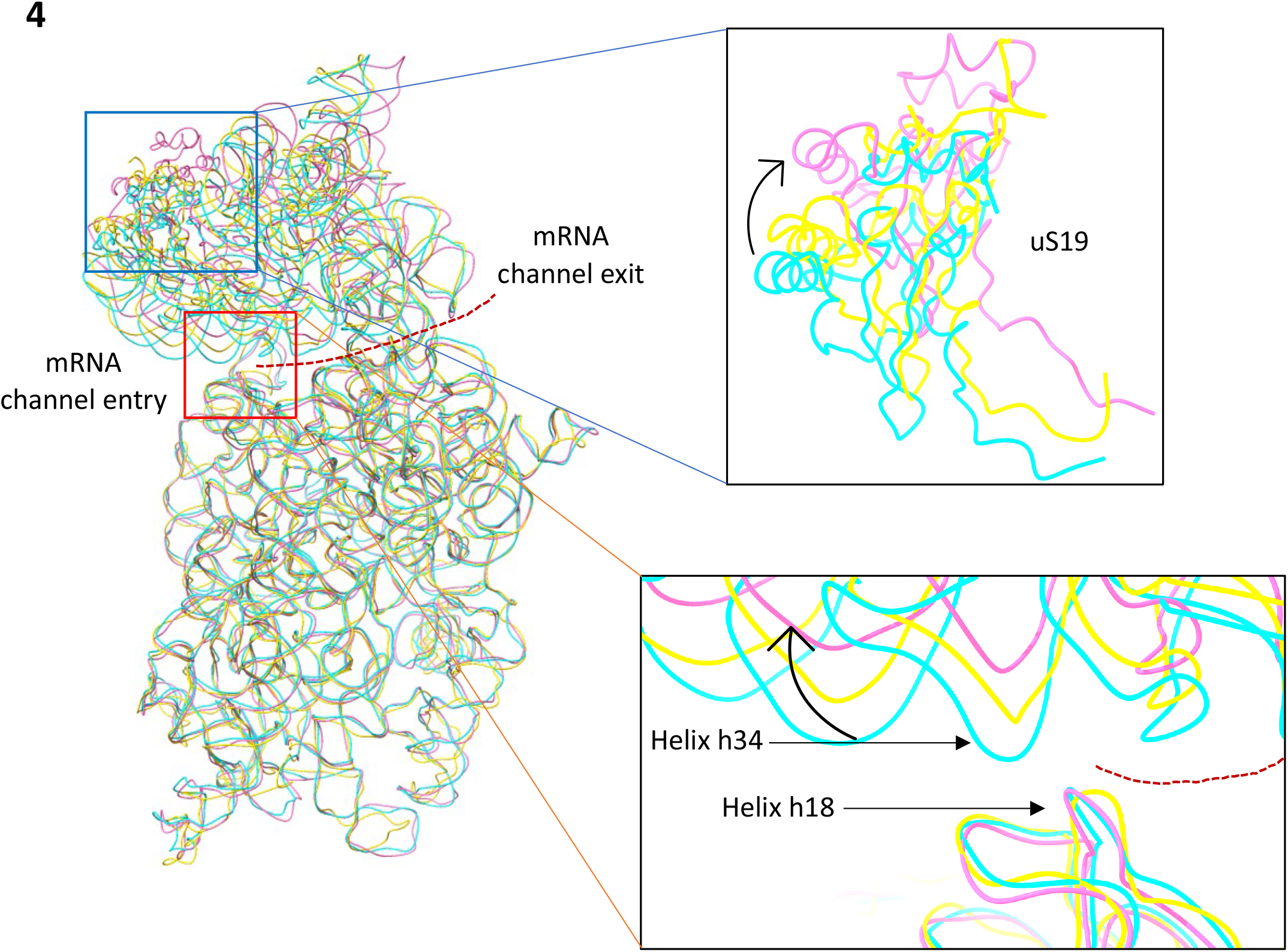
eIF4B stabilizes the partially open state of the 40S head. Comparison of the 40S mRNA channel latch conformations (PDB: 6GSM-open conformation in pink, 6GSN-closed conformation in cyan, and 40S-eIF4B-partially open conformation in yellow). Superimposition of the coordinates shows movement in the 40S head. Helices h18 and h34 form the latch and the curved arrow shows the direction of 40S head movement with it being in a partially rotated state and hence in a partially open conformation in 40S-eIF4B (yellow). The movement of the head can also be observed by the rotation of the proteins present in the 40S head like uS19.

The7ocationn of eIF4B on the 40S in the 40S-eIF4B map clashes with the RRM domain of eIF3g in the context of the yeast 43S PIC (Figure 5A) (Kratzat *et al*., 2021). eIF3 is a large multi-subunit complex composed of 12 (a-m) subunits in mammals and 6 (a, b, c, g, i, and j) subunits in *S. cerevisiae*. The subunits eIF3 a and c have the PCI/MPN domains and form the core eIF3 unit present at the mRNA exit channel of the ribosome. eIF3b, eIF3g, and eIF3i form the flexible peripheral core of the eIF3 complex which is reported to relocate from the solvent-exposed to the inter-subunit interface of the 40S during mRNA recruitment (Llacer *et al*., 2021). eIF3b and eIF3i have a nine and seven-bladed β-propeller and eIF3b and eIF3g have an RRM domain in the N- and C-terminus, respectively. The cryo-EM structures reveal that the NTD of eIF3g is sandwiched between the 40S body and β-propeller domain of eIF3i whereas its C-terminal RRM domain is present at the mRNA entry channel (Kratzat *et al*., 2021). Biochemical studies such as pull-downs and far western experiments have revealed that eIF3g-RRM can directly interact with the ribosomal proteins uS3 and uS10 (Cuchalová *et al*., 2010). This makes the presence of eIF3g-RRM and eIF4B at the mRNA channel entry mutually exclusive as they are occupying and interacting with the same protein (uS3) present there (Figure 5A). As mentioned above, the eIF3b-g-i complex is present on the solvent-exposed side of the 43S PIC (Kratzat *et al*., 2021) and relocates to the inter-subunit interface in the 48S PIC when the mRNA is recruited (Llacer *et al*., 2021). It is likely that the displacement of eIF3g-RRM by eIF4B contributes to the relocation of the entire eIF3b-g-i complex to the inter-subunit interface of the 40S.

**Figure 5:**
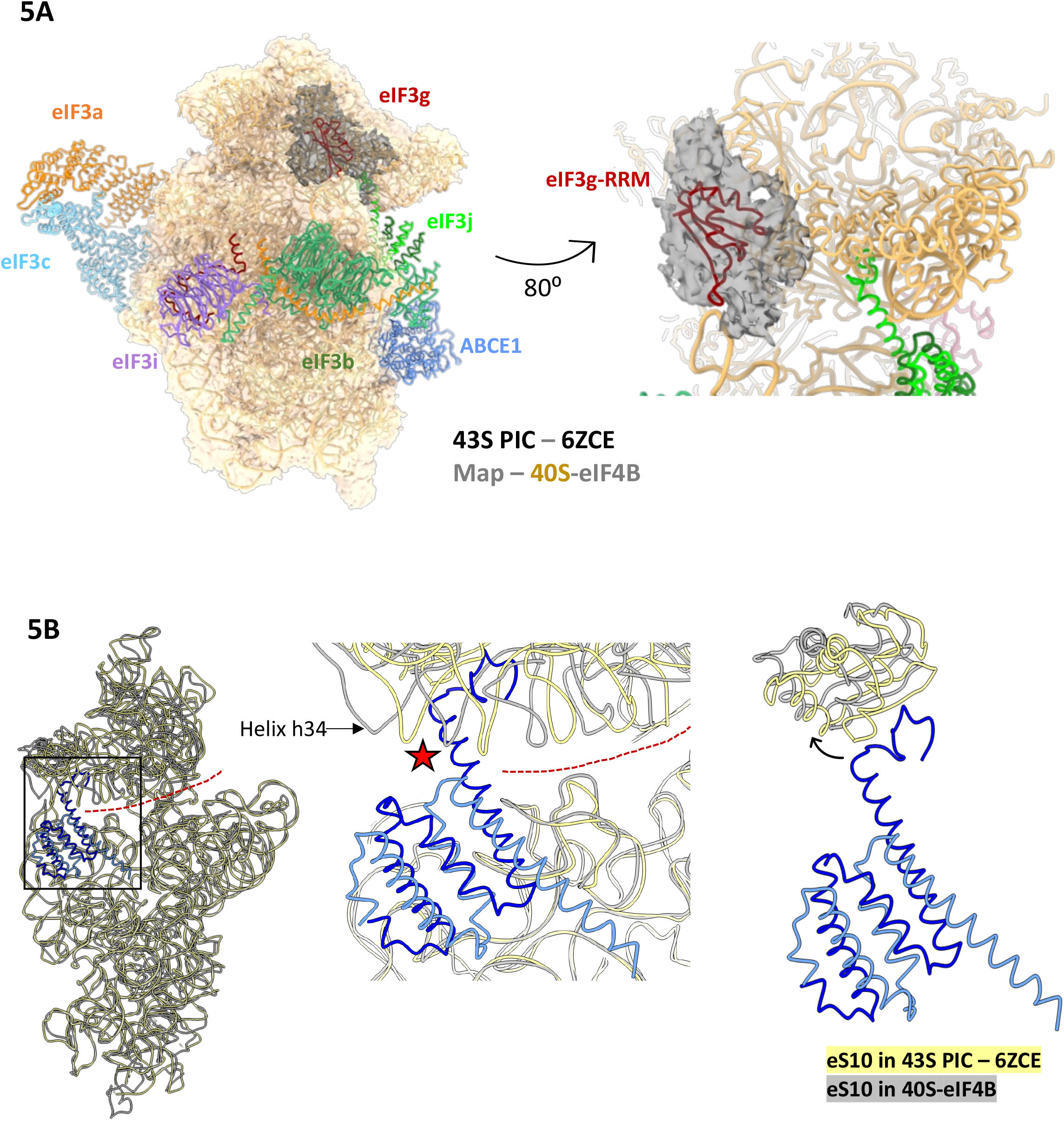
Potential consequences of the 40S-eIF4B interaction in the context of the 43S PIC. (A) Superimposition of the native 43S PIC from *S. cerevisiae* with eIF3 subunits present on the solvent-exposed side in the 40S-eIF4B map (yellow – 40S, gray – eIF4B) shows that the eIF3g-RRM domain (red) occupies the eIF4B density completely. This suggests that in the context of the 48S PIC, there exists a mutually exclusive relation of the eIF3g-RRM domain and eIF4B at this location. (B) eIF3j binds and blocks the mRNA channel of 40S (dashed red line) in the context of the 43S PIC (PDB: 6ZCE - yellow. eIF3j dimer – dark and light blue). The C-terminal tail of eIF3j has a severe steric clash with rRNA helix h34 when the 40S mRNA channel latch is in a partially open state in 40S-eIF4B (gray). eS10, which interacts with eIF3j-CTT moves from its original location in the 43S PIC (yellow) due to the opening of the mRNA channel latch in 40S-eIF4B (gray).

**Figure 6:**
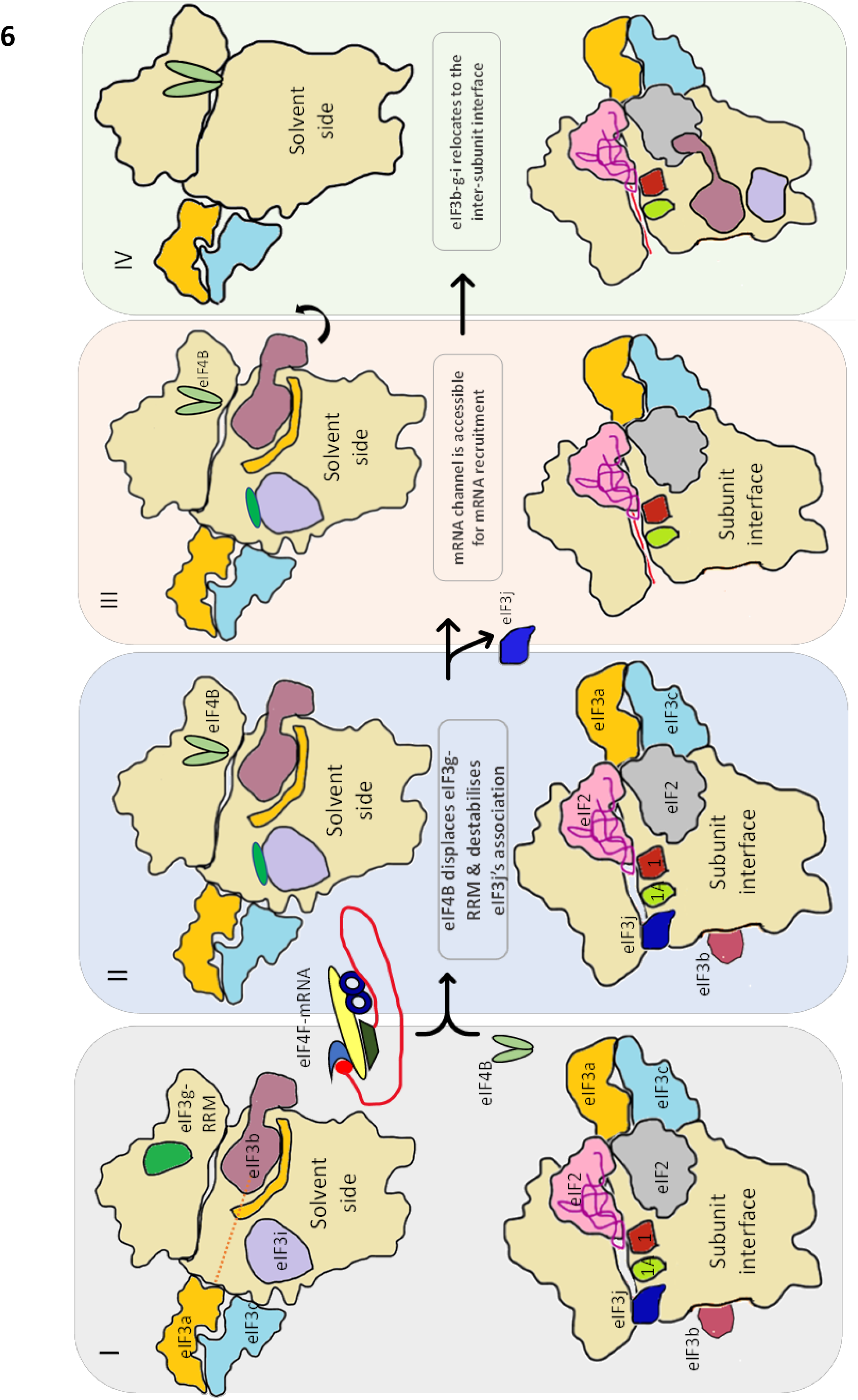
(I-IV) Proposed model explaining the events that might occur during translation initiation in yeast. In the 43S PIC, eIF3g-RRM is located on the solvent side near the mRNA channel entry on the 40S (I). When eIF4B comes either in association with eIF4F-mRNA or alone, it displaces eIF3g-RRM and occupies the same position as that of eIF3g-RRM (II). eIF4B’s association may also destabilize eIF3j’s interaction with the 40S due to competition for interaction with the same ribosomal proteins. This may lead to the dissociation of eIF3j, which removes the blockage it creates for mRNA’s entry and accommodation in the mRNA channel of the 40S (III). The displacement of eIF3g-RRM by eIF4B further triggers the relocation of the eIF3b-g-i peripheral core to the 40S inter-subunit interface where it helps in start codon selection (IV).

Along with eIF3g, yeast eIF4B may also affect the interaction of eIF3j with the 40S. eIF3j is an accessory and sub-stoichiometric factor playing important role in mRNA quality control, preventing premature recruitment of mRNA and association of the 60S subunit (Hinnebusch, 2017). Hydroxy radical cleavage (Fraser *et al*., 2007) and cryo-EM map of native 43S PIC (Kratzat *et al*., 2021) have placed eIF3j such that its C-terminal tail region (CTT) occupies and blocks the mRNA channel of the 40S by interacting with h34). Therefore, for mRNA to occupy the 40S mRNA channel, eIF3j must dissociate. The decrease in the affinity of human eIF3j for the 40S in the presence of mRNA and eIF4 factors (Sokabe and Fraser, 2017), the absence of eIF3j in the native yeast 48S PIC structure (Kratzat *et al*., 2021) (PDB id: 6ZU9), suggest that eIF4 factors may have a role in the dissociation of eIF3j. When we overlapped the coordinates of the eIF3j-bound 43S PIC (PDB id: 6ZCE) with those of the 40S coordinates refined in the 40S-eIF4B map, it was observed that eIF3j-CTT clashes with h34 in 40S-eIF4B (Figure 5B). Further, eIF3j-CTT interacts with eS10 in the 43S PIC. However, we observe that due to the movement of the 40S-head in the 40S-eIF4B structure with respect to the head conformation in the closed mRNA latch state, eS10 present in the 40S head moves away from eIF3j-CTT and the interaction between them is likely to break (Figure 5B). Thus, the clash with h34 and destabilized interaction with eS10 may explain the decrease in eIF3j’s affinity for the 40S in the presence of eIF4 factors (Sokabe and Fraser, 2017).

eIF4 factors, mainly eIF4G and eIF4E, are targeted by viruses in multiple ways to hijack host translational machinery (Jaafar and Kieft, 2019). Interestingly, the position on the 40S occupied by eIF4B and the eIF3g-RRM domain is also the binding site for the SARS-CoV-2 RNA encoded protein Nsp1 (PDB – 6ZLW, 6ZOJ; Figure S4B). The N-terminal domain of Nsp1 binds the 40S near the mRNA entry channel with its C-terminal helices present in the mRNA channel, inhibiting host mRNA accommodation recruitment and ceasing host translation (Schubert *et al*., 2020; Thoms *et al*., 2020). Drug targeting the C-terminal helices of Nsp1 prevents the shutdown of host translation (Afsar *et al*., 2022). The shared binding site of Nsp1 with eIF4B and eIF3g-RRM hints that the viral protein Nsp1 competes with host eIFs for binding to the 40S (Figure S4B).

### Concluding remarks

The map of yeast 40S-eIF4B highlights the position and binding partners of eIF4B on the 40S ribosomal subunit along with the possible ways in which eIF4B may help in the mRNA recruitment during translation initiation. Its presence near the mRNA entry channel of the 40S suggests that eIF4B helps stabilize the open mRNA latch of 40S to facilitate mRNA recruitment. Several biochemical and genetics-based studies further support this role of eIF4B (Sen *et al*., 2016; Walker *et al*., 2013; Zhou *et al*., 2014). The same binding site of eIF4B with the eIF3g-RRM domain at the mRNA entry site hints that the signal for the relocation of the eIF3b-g-i complex could be the binding of eIF4B. Along with this, eIF4B could also be responsible for the removal of eIF3j-caused blockage of the 40S mRNA channel by facilitating its dissociation. Based on our observations and comparison of 40S-eIF4B with reported structures of partial yeast 43S and 48S PICs, were have proposed a model explaining the events that may happen during mRNA recruitment to the small ribosomal subunit in yeast (Figure 5). While we were finalizing this manuscript a structure of human 48S PIC was reported where a low-resolution density in the vicinity of the mRNA channel entry on the solvent-exposed side is tentatively assigned to eIF4B (Brito Querido *et al*., 2022). Overall, the structural analysis presented in this study shows how yeast eIF4B helps in mRNA recruitment as well as scanning during translation initiation.

## Methods

### Purification of 40S ribosomal subunit

The 40S ribosomal subunit was purified using earlier mentioned protocols (Acker *et al*., 2007; Fernandez *et al*., 2014) with some modifications. *K. lactis* pellet from 6l was resuspended in 1X ribo-lysis buffer (3X the pellet volume, 20 mM MES pH 6.0, 150 mM Potassium acetate pH 7.6, 10 mM Magnesium acetate, 0.2 mM PMSF, 0.2 mM benzamidine, 2 mM DTT, protease inhibitor tablet). The popcorns were lysed using a blender and lysate was clarified by centrifugation at 13000 rpm for 30 minutes at 4°C. 22.5 ml lysate was layered on 2.5 ml sucrose cushion (1M sucrose in ribo-lysis buffer) at centrifuged at 28000 rpm for 6 hours at 4°C. The 80S pellet was washed with ice-cold high salt wash buffer and resuspended in the high salt wash buffer (20 mM MES pH6.0, 100 mM potassium acetate, 2.5 mM magnesium acetate, 500 mM KCl, 0.2 mM PMSF, 0.2 mM benzamidine, 2 mM DTT, 1 mg/ml heparin). The resuspended 80S was layered on a sucrose cushion made in high salt buffer and centrifuged at 45000 rpm for 8 hours at 4°C. The 80S pellet was resuspended in dissociation buffer (20 mM MES pH6.0, 600 mM KCl, 8 mM magnesium acetate, 0.2 mM PMSF, 0.2 mM benzamidine, 2 mM DTT, 1 mg/ml heparin) such that the final OD at 260 nm is 100-150. 1 ml 80S was layered on 10-35% sucrose gradient (20 mM MES pH 6.0, 600 mM KCl, 8 mM magnesium acetate, 0.2 mM PMSF, 0.2 mM benzamidine, 2 mM DTT, 1 mg/ml heparin, 10% or 35% sucrose) after 1 mM puromycin treatment for 1 hour at 4°C. The gradient was centrifuges at 25000 rpm for 8 hours at 4°C. 1ml fractions were collected and analysed on 1% agarose gel to check for fractions containing 40S. 40S containing fractions were pooled and diluted 3X with dissociation buffer and layered on 1 M sucrose cushion made in ribosome storage buffer (20 mM HEPES pH 7.4, 100 mM potassium acetate pH 7.6, 2.5 mM magnesium acetate, 250 mM sucrose, 2 mM DTT) followed by centrifugation at 52000 rpm for 8 hours at 4°C. The 40S pellet was resuspended in ribosome storage buffer and further concentrated if required using the viva-spin concentrator (100kDa cut-off).

### Purification of eIF4B

eIF4B was purified as mentioned by Liu *et al*., 2019 followed by size exclusion chromatography using the HiLoad 200pg column.

### Reconstitution of the 40S-eIF4B complex and data collection

240nM 40S was incubated at room temperature (25°C) for 5 minutes followed by the addition of 10-fold molar excess of eIF4B. The complex was incubated at room temperature for 5 minutes and 3ul was applied to a glow discharged Quantifoil R2/2 grid. Data was collected on Thermo Scientific Talos Arctica operated at 200 kV under low dose conditions (60 e^-^/Å^2^) using a defocus range of 1.5-3µm. Images were recorded on K2 detector with a magnification of 42K (pixel size – 1.17Å).

### Cryo-EM data processing

Data processing was done using RELION 3.1.1 (Figure S1). After the motion-correction and CTF-correction of 7,975 movie files, LoG-based auto picking was performed, resulting in 1,492,547 particles. After multiple rounds of 2D and 3D classification to remove junk particles, 593,832 good particles were used for 3D refinement (Figure S2A). During initial 3D classification runs we observed extra density near mRNA entry channel in one class. Hence, we decided to perform the 3D focussed classification of the particles into two classes with a mask focused on the mRNA channel entry region of the 40S ribosome on the solvent-exposed side. Class 1 and 2 had 108,616 and 465,216 particles, respectively. These classes were individually subjected to Bayesian polishing, CTF refinement, 3D auto-refinement, and post-processing, which resulted in two maps of 4.6Å and 4Å global resolution. The resolution was calculated as per the FSC=0.143 cutoff.

### Model building and Refinement

The atomic model of the head and body of *K. lactis* 40S (PDB ID: 3JAM) was initially rigid body fit separately in Chimera (Pettersen *et al*., 2004) and they were real space refined separately in Phenix (Emsley *et al*., 2010). The real-space refined models of 40S head and body were merged in Coot to get the model of 40S. All figures were generated using ChimeraX (Pettersen *et al*., 2021).

Accession Codes: Maps of the 40S and 40S-eIF4B have been deposited in the EMDB with accession codes EMD-XXXX and EMD-XXXX. Atomic coordinates of the partially open state of 40S corresponding to the cryo-EM map of 40S-eIF4B have been deposited in the PDB with accession code XXXX.

## Acknowledgements

We are thankful to the IISc cryo-EM facility for grid preparation and data collection. We thank members of the TH lab and Sarah Walker for the insightful discussion. This work was supported by Intermediate Fellowship from DBT-Welcome Trust India Alliance to TH (IA/I/17/2/503313).

## Supplementary Figures

**Figure S1: Cryo-EM data processing pipeline used for processing the 40S-eIF4B dataset**

**Figure S2: 2D classes, angular distribution plot and FSC curves**

(A) The clean 2D classes comprising 573,832 particles were used as the input for 3D classification. (B) The angular distribution plot of the 40S-eIF4B auto-refined map shows that majority of the views are represented in this class (shown by red and blue bars). However, some views are still missing (white spaces) and this is due to the preferential orientation of the 40S on the cryo-EM grid. (C and D) FSC curves of the 40S and 40S-eIF4B gives the global resolution of the maps at FSC 0.143 cut-off.

**Figure S3: eIF4B density in the 40S-eIF4B map**

(A and B) The extra density of eIF4B at different thresholds (left and right panels) of the (A) post-processed map and (B) low pass filtered map shows the density originating near uS3 and rRNA helix h16. (C) Local resolution of the 40S-eIF4B map. The resolution of the core of the 40S body is higher (around 4Å) when compared to its peripheral regions as well as the 40S head. (right hand panel) Resolution of the extra density tentatively assigned to eIF4B ranges from 8-10Å which makes model building difficult.

**Figure S4: Binding site of eIF4B and SARS-CoV-2 NSP1 on the 40S overlaps**

(A) The RRM domain of human eIF4B (blue; PDB – 6FEC) completely fits into the extra density tentatively assigned to the yeast eIF4B in our 40S-eIF4B map. (B) SARS-CoV-2 NSP1 (EMD11276; NSP1-cyan, mammalian 40S-yellow) binds and the 40S at the same site at which yeast eIF4B (yeast eIF4B-purple, yeast 40S-pink) binds during mRNA recruitment. Therefore, by blocking this site, NSP1 might prevent the host mRNA recruitment, thus hijacking the host translation system.

## Legends for Movies

**Movie 1: Complete 360° rotation of the 40S-eIF4B cryo-EM map**

The refined coordinates of 40S are shown and the extra density at the mRNA entry site on the solvent face is shown in magenta colour.

## References

Acker MG, Kolitz SE, Mitchell SF, Nanda JS, Lorsch JR (2007) Reconstitution of yeast translation initiation. Methods Enzymol. 430: 111–145

Afsar M, Narayan R, Akhtar M N, Das D, Rahil H, Nagaraj S K, and Hussain T (2022). Drug targeting Nsp1-ribosomal complex shows antiviral activity against SARS-CoV-2. Elife, 11, e74877.

Aitken CE, Lorsch JR (2012) A mechanistic overview of translation initiation in eukaryotes. Nat. Struct. Mol. Biol.19: 568–576

Altmann M, Muller PP, Wittmer B, Ruchti F, Lanker S, Trachsel H (1993) A Saccharomyces cerevisiae homologue of mammalian translation initiation factor 4B contributes to RNA helicase activity. EMBO J 12: 3997–4003

Andreou AZ, Klostermeier D (2014) eIF4B and eIF4G jointly stimulate eIF4A ATPase and unwinding activities by modulation of the eIF4A conformational cycle. J Mol Biol 426: 51–61

Andreou AZ, Harms U, and Klostermeier D (2017). eIF4B stimulates eIF4A ATPase and unwinding activities by direct interaction through its 7-repeats region. RNA Biol., 14(1): 113–123.

Aylett C H and Ban N (2017). Eukaryotic aspects of translation initiation brought into focus. Philos. Trans. R. Soc. Lond., B, Biol. Sci., 372(1716), 20160186.

Brito Querido J, Sokabe M, Kraatz S, Gordiyenko Y, Skehel JM, Fraser CS, Ramakrishnan V (2020) Structure of a human 48S translational initiation complex. Science 369: 1220–1227

Brito Querido J, Sokabe M., Díaz-López I, Gordiyenko Y, Fraser CS and Ramakrishnan V (2022) The structure of a human translation initiation complex reveals two independent roles for the helicase eIF4A. bioRxiv.

Cuchalová, L., Kouba, T., Herrmannová, A., Dányi, I., Chiu, W. L., and Valásêk, L. (2010). The RNA recognition motif of eukaryotic translation initiation factor 3g (eIF3g) is required for resumption of scanning of posttermination ribosomes for reinitiation on GCN4 and together with eIF3i stimulates linear scanning. Mol. Cell. Biol., 30(19), 4671–4686.

Eliseev B, Yeramala L, Leitner A, Karuppasamy M, Raimondeau E, Huard K, Alkalaeva E, Aebersold R, Schaffitzel C (2018) Structure of a human cap-dependent 48S translation pre-initiation complex. Nucleic Acids Res. 46: 2678–2689

Emsley P, Lohkamp B, Scott WG, Cowtan K (2010) Features and development of Coot. Acta Crystallogr., Sect D 66: 486–501

Fernandez IS, Bai XC, Murshudov G, Scheres SH, Ramakrishnan V (2014) Initiation of translation by cricket paralysis virus IRES requires its translocation in the ribosome. Cell 157: 823–831

Fraser CS (2015) Quantitative studies of mRNA recruitment to the eukaryotic ribosome. Biochimie 114: 58–71

Fraser CS, Berry KE, Hershey JW, Doudna JA (2007) eIF3j is located in the decoding center of the human 40S ribosomal subunit. Mol Cell 26: 811–819

Hernandez G, Vazquez-Pianzola P (2005) Functional diversity of the eukaryotic translation initiation factors belonging to eIF4 families. Mech Dev 122: 865–876

Hinnebusch AG (2017) Structural Insights into the Mechanism of Scanning and Start Codon Recognition in Eukaryotic Translation Initiation. Trends Biochem Sci 42: 589–611

Jaafar ZA, Kieft JS (2019) Viral RNA structure-based strategies to manipulate translation. Nat. Rev. Microbiol. 17: 110–123

Jackson RJ, Hellen CU, Pestova TV (2010) The mechanism of eukaryotic translation initiation and principles of its regulation. Nat Rev Mol Cell Biol 11: 113–127

Kozak M (1986) Point mutations define a sequence flanking the AUG initiator codon that modulates translation by eukaryotic ribosomes. Cell 44: 283–292

Kratzat H, Mackens-Kiani T, Ameismeier M, Potocnjak M, Cheng J, Dacheux E, Namane A, Berninghausen O, Herzog F, Fromont-Racine M et al (2021) A structural inventory of native ribosomal ABCE1-43S pre-initiation complexes. EMBO J 40: e105179

Li Z, Cheng Z, Raghothama C, Cui Z, Liu K, Li X, Jiang C, Jiang W, Tan M, Ni X et al (2018) USP9X controls translation efficiency via deubiquitination of eukaryotic translation initiation factor 4A1. Nucleic Acids Res.46: 823–839

Liu X, Moshiri H, He Q, Sahoo A, Walker SE (2021) Deletion of the N-Terminal Domain of Yeast Eukaryotic Initiation Factor 4B Reprograms Translation and Reduces Growth in Urea. Front Mol Biosci 8: 787781

Liu X, Schuessler PJ, Sahoo A, Walker SE (2019) Reconstitution and analyses of RNA interactions with eukaryotic translation initiation factors and ribosomal preinitiation complexes. Methods 162-163: 42–53

Llacer JL, Hussain T, Dong J, Villamayor L, Gordiyenko Y, Hinnebusch AG (2021) Large-scale movement of eIF3 domains during translation initiation modulate start codon selection. Nucleic Acids Res.49: 11491–11511

Llacer JL, Hussain T, Marler L, Aitken CE, Thakur A, Lorsch JR, Hinnebusch AG, Ramakrishnan V (2015) Conformational Differences between Open and Closed States of the Eukaryotic Translation Initiation Complex. Mol Cell 59: 399–412

Mayberry LK, Allen ML, Dennis MD, Browning KS (2009) Evidence for variation in the optimal translation initiation complex: plant eIF4B, eIF4F, and eIF(iso)4F differentially promote translation of mRNAs. Plant physiology 150: 1844–1854

Merrick WC (2015) eIF4F: a retrospective. J. Biol. Chem. 290: 24091–24099

Mishra RK, Datey A, Hussain T (2020) mRNA Recruiting eIF4 Factors Involved in Protein Synthesis and Its Regulation. Biochemistry 59: 34–46

Pettersen EF, Goddard TD, Huang CC, Couch GS, Greenblatt DM, Meng EC, Ferrin TE (2004) UCSF Chimera--a visualization system for exploratory research and analysis. J. Comput. Chem. 25: 1605–1612

Pettersen EF, Goddard TD, Huang CC, Meng EC, Couch GS, Croll TI, Morris JH, Ferrin TE (2021) UCSF ChimeraX: Structure visualization for researchers, educators, and developers. Protein Sci. 30: 70–82

Rogers GW, Jr., Richter NJ, Merrick WC (1999) Biochemical and kinetic characterization of the RNA helicase activity of eukaryotic initiation factor 4A. J. Biol. Chem. 274: 12236–12244

Rossler I, Embacher J, Pillet B, Murat G, Liesinger L, Hafner J, Unterluggauer JJ, Birner-Gruenberger R, Kressler D, Pertschy B (2019) Tsr4 and Nap1, two novel members of the ribosomal protein chaperOME. Nucleic Acids Res.47: 6984–7002

Schubert K, Karousis ED, Jomaa A, Scaiola A, Echeverria B, Gurzeler LA, Leibundgut M, Thiel V, Muhlemann O, Ban N (2020) SARS-CoV-2 Nsp1 binds the ribosomal mRNA channel to inhibit translation. Nat. Struct. Mol. Biol. 27: 959–966

Sen ND, Zhou F, Harris MS, Ingolia NT, Hinnebusch AG (2016) eIF4B stimulates translation of long mRNAs with structured 5’ UTRs and low closed-loop potential but weak dependence on eIF4G. Proc Natl Acad Sci U.S.A. 113: 10464–10472

Shahbazian D, Parsyan A, Petroulakis E, Topisirovic I, Martineau Y, Gibbs BF, Svitkin Y, Sonenberg N (2010) Control of cell survival and proliferation by mammalian eukaryotic initiation factor 4B. Mol Cell Biol 30: 1478–1485

Sokabe M, Fraser CS (2017) A helicase-independent activity of eIF4A in promoting mRNA recruitment to the human ribosome. Proc Natl Acad Sci U.S.A. 114: 6304–6309

Thoms M, Buschauer R, Ameismeier M, Koepke L, Denk T, Hirschenberger M, Kratzat H, Hayn M, Mackens-Kiani T, Cheng J et al (2020) Structural basis for translational shutdown and immune evasion by the Nsp1 protein of SARS-CoV-2. Science 369: 1249–1255

Uppala JK, Sathe L, Chakraborty A, Bhattacharjee S, Pulvino AT, Dey M (2022) The cap-proximal RNA secondary structure inhibits preinitiation complex formation on HAC1 mRNA. J. Biol. Chem.298: 101648

Walker SE, Zhou F, Mitchell SF, Larson VS, Valasek L, Hinnebusch AG, Lorsch JR (2013) Yeast eIF4B binds to the head of the 40S ribosomal subunit and promotes mRNA recruitment through its N-terminal and internal repeat domains. RNA 19: 191–207

Zhou F, Walker SE, Mitchell SF, Lorsch JR, Hinnebusch AG (2014) Identification and characterization of functionally critical, conserved motifs in the internal repeats and N-terminal domain of yeast translation initiation factor 4B (yeIF4B). J. Biol. Chem.289: 1704–1722

