## Supplementary Figures for "Yeast eukaryotic initiation factor 4B remodels the mRNA entry site on the small ribosomal subunit"

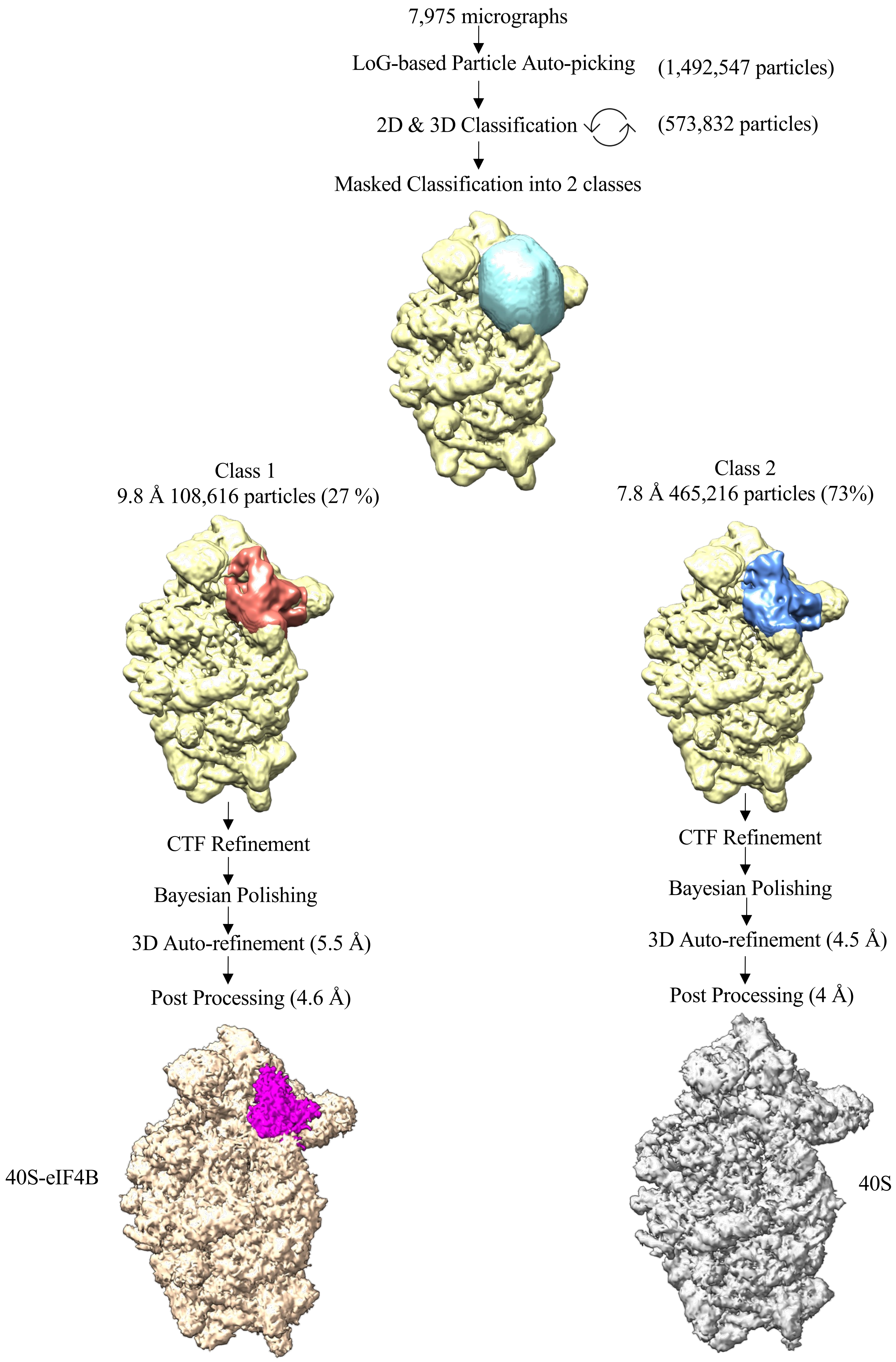


**Figure S1: Cryo-EM data processing pipeline of 40S-eIF4B**

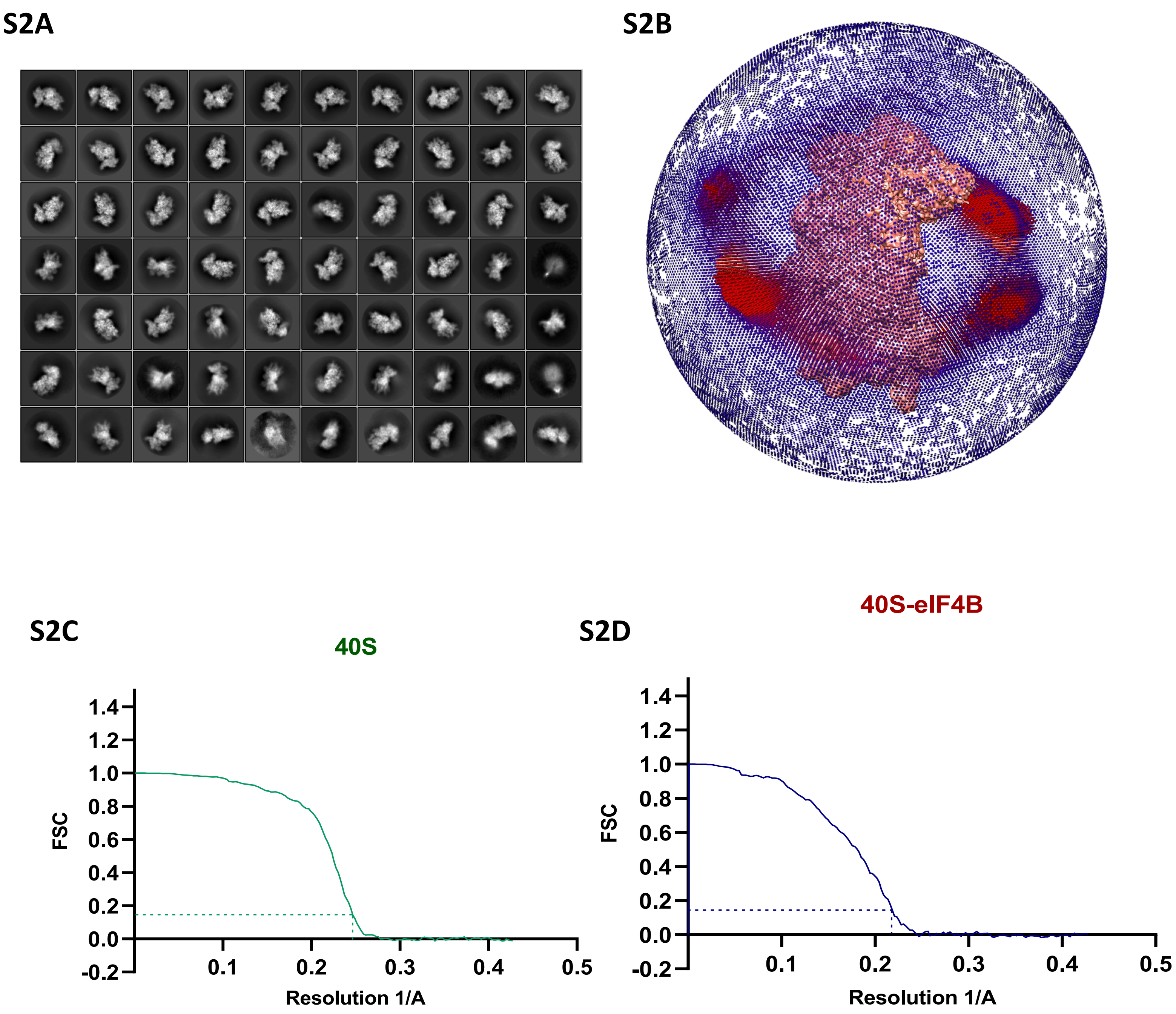


**Figure S2: 2D classes, angular distribution plot and FSC curves**

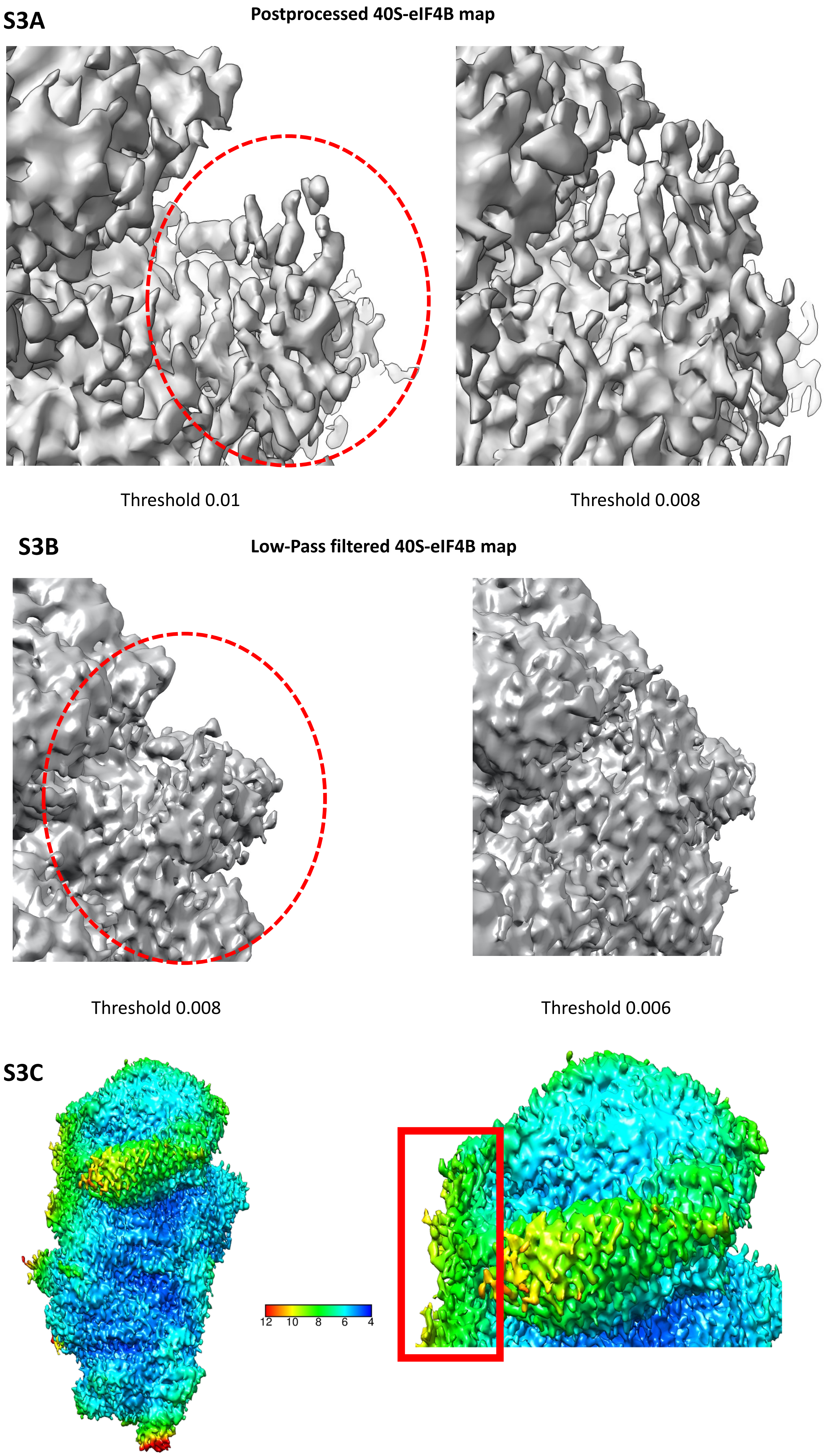


**Figure S3: eIF4B density in the 40S-eIF4B map**

S4A

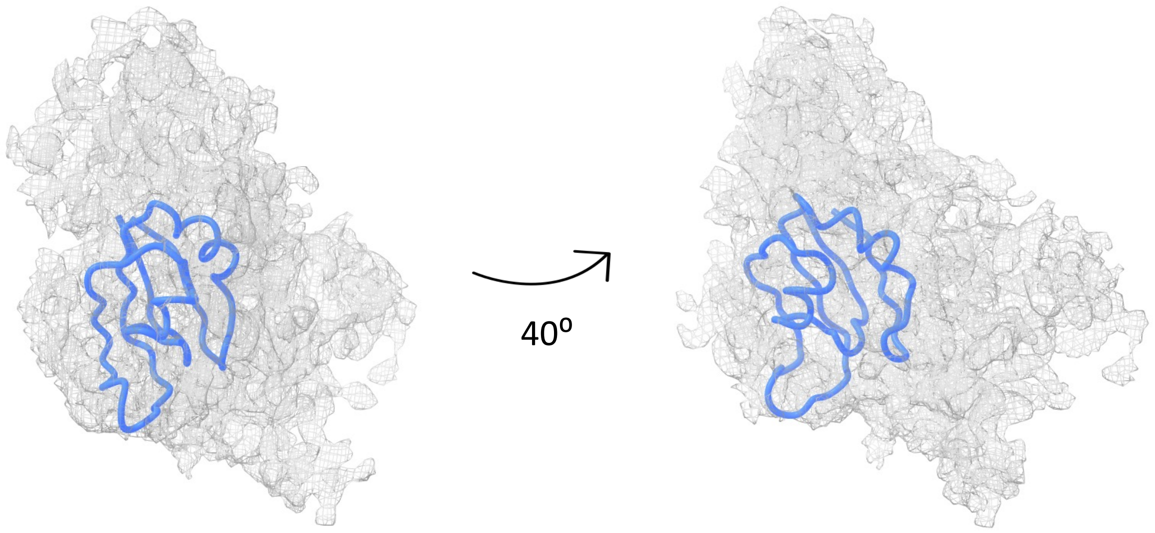


S4B

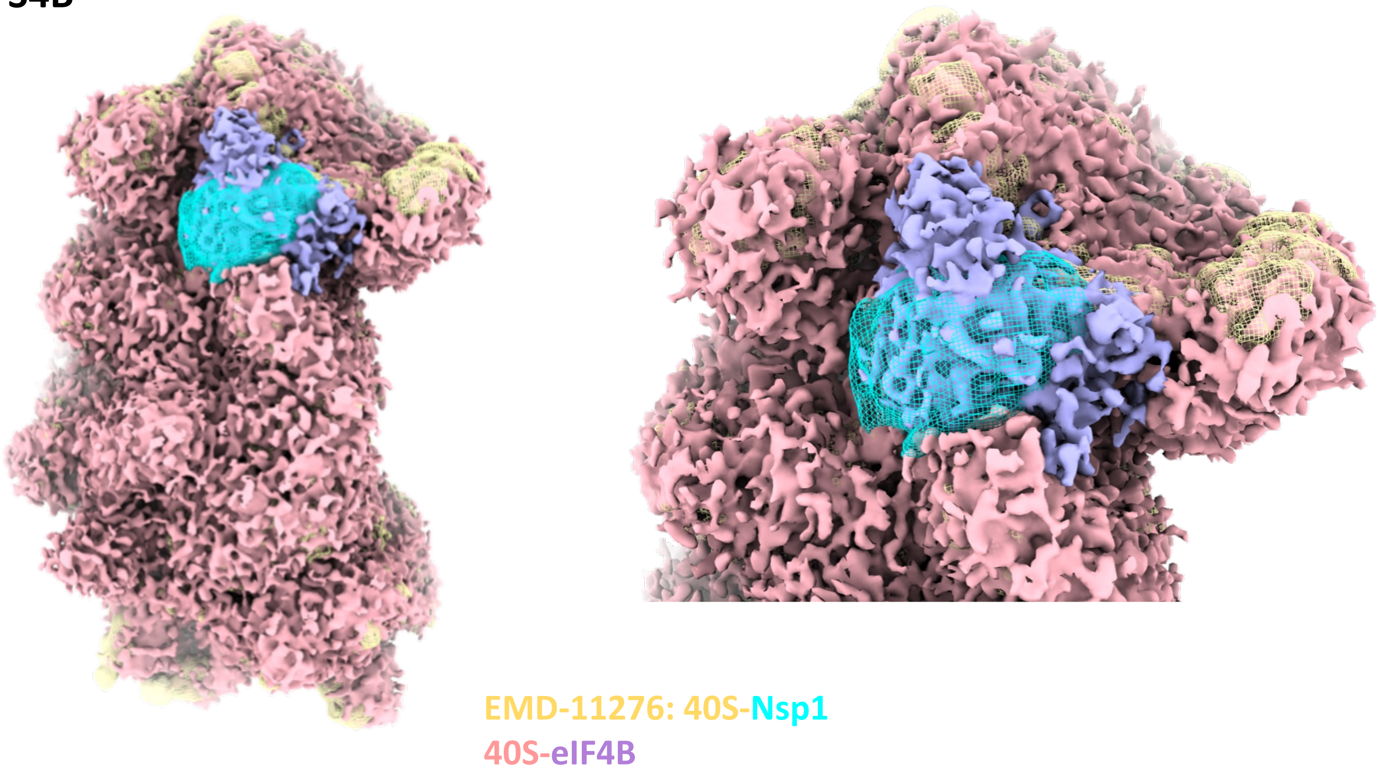


**Figure S4: Binding site of eIF4B and SARS-CoV-2 NSP1 on the 40S overlaps**
